# Trimethoprim suppress phage activity and reduces toxin production in a clinical STEC strain, even in the presence of DNA-damaging antibiotics

**DOI:** 10.64898/2026.09.17.752489

**Authors:** Ayesha Umair, Alexander P. Hynes

**Affiliations:** Department of Medicine, McMaster University; Department of Biochemistry and Biomedical Sciences, McMaster University

## Abstract

Shiga toxin-producing *Escherichia coli* (STEC) are virulent due to their production of lethal toxins, which are genetically encoded by viruses that exclusively infect these bacteria, called phages. After infection, children and immunocompromised patients are particularly at risk of developing life-threatening complications. Antibiotics are contraindicated here, because they increase phage activity, and this leads to increased toxin production. This limits treatments to fluid resuscitation and supportive care.

Here, we describe how acid exposure and trimethoprim both potently suppress phage production ∼100-fold, even in the presence of antibiotics that increase phage activity. We characterized these stimuli further, and found that they suppress phages independently of canonical bacterial SOS responses that are well-established to be involved in phage activity. Trimethoprim partially relies on the glutamate-dependent bacterial acid stress response to have this suppressive effect. It allows the earliest steps of phage induction to happen, but completely blocks their genomic replication. We translated these findings to a clinical STEC isolate and found that trimethoprim potently suppressed toxin production, again, even in the presence of antibiotics that increase phage activity. These findings form a key lead in finding a new option to treat these infections, without risking potentially lethal complications.

## Main

Bacteriophages, or viruses that infect bacteria, are often responsible for encoding virulence factors like toxins or defense mechanisms that increase the fitness of their hosts.^1^ Through the presence of these virulence factors, bacteria can become more dangerous as pathogens against humans.^1^ Shiga-toxin encoding *Escherichia coli* (STEC) are a notable example; they are infected by Stx phages, which are a group of morphologically diverse phages that encode Shiga toxins Stx1 and Stx2.^2^

STEC infections can be life-threatening. Up to 20% of children with high-risk STEC infections (usually defined as encoding Stx2) will develop hemolytic uremic syndrome (HUS), a dangerous complication defined by the breakdown of erythrocytes, platelet depletion, and kidney damage.^3^ Severe acute kidney injury occurs in one-half of cases, and as many as one-half to two-thirds of patients with HUS require dialysis.^4^ The toxins are responsible for HUS; they enter the bloodstream, causing end-organ damage through protein synthesis inhibition and apoptosis.^5^ HUS, in turn, can lead to important long-term sequelae: 36% of patients presented with hypertension and impaired kidney function over a year after developing HUS.^6^

The production of Shiga toxins is strongly linked to phage activity.^7^ When these phages undergo a lytic cycle, they hijack the host cell machinery to produce viral progeny, and then lyse the cell to release them. Toxin production occurs late in this cycle, and release is mediated by phage lysis.^7^ Because Stx phages are temperate phages, they can also undergo a different “lysogenic” life cycle, where the phage genetic material integrates into the host genome and is replicated alongside the host as a prophage. The bacterium carrying the largely quiescent phage is called a lysogen, and, by virtue of carrying the phage, the lysogen now encodes the Shiga toxin. When the lysogen encounters a stressor, canonically a DNA-damaging stimulus such as UV light, reactive oxygen species, or DNA-damaging antibiotics, the prophage goes from the lysogenic cycle to the phage lytic cycle in a process known as induction. Accordingly, phage induction leads to increased toxin production and release.^8^

Since antibiotics can cause phage induction and increase toxin production, their use is contraindicated in STEC infections because of this relationship between toxins and phages. Administering antibiotics during STEC infections can increase the risk of HUS by ∼20-fold.^9^ Currently, clinical guidelines focus primarily on hydration and supportive management for these infections.^10^ Azithromycin, a protein synthesis inhibitor, was explored in preclinical and clinical work. It was found to limit toxin production *in vitro*.^11^ Gnotobiotic piglets infected with STEC and treated with azithromycin showed reduced intestinal toxin levels, and recovered with little or no complications.^12^ It was also the focus of a clinical trial aimed at treating STEC-HUS^13^ but results have not been published; other studies showed that azithromycin use still increased the risk of HUS.^14^ Concerns also exist around the azithromycin’s arrhythmogenic potential, especially in patients with the electrolyte imbalances from kidney injury in HUS.^15^

Instead of approaches that limit toxin production, we focus on the underlying phage activity. If phage induction could be suppressed, it could lead to better, more potent upstream treatment options for STEC infections. Researchers have tried suppressing phages in the context of STEC *in vitro*. Apyrase, which cleaves and neutralizes ATP, and nitric oxide are both known to decrease Stx2 levels.^16,17^ Others have discussed Stx phages being suppressed through activation of the general RpoS stress response.^18^

Here, we present acute acid exposure as a mechanism for the potent suppression of phages, even in the presence of DNA-damaging drugs. We also see this effect translated to trimethoprim overriding toxin suppression in a clinical STEC isolate. These results could be valuable in treating STEC infections, and also in exploring a key pathway in bacteria at the center of acid stress responses, DNA damage, and phage induction.

### Acute acid exposure to pH 5 leads to a potent decrease in phage titers

We explored the effect of various environmental conditions to see if they could lead to phage suppression. Lysogens and phages are known to encounter stimuli in their natural surroundings that influence baseline spontaneous induction in the absence of DNA-damaging stimuli, such as salt stress and compounds released by competing bacteria.^19,20^ Nonpathogenic, laboratory strains of *E. coli* lysogens harboring model temperate phage HK97 were grown to the beginning of the exponential phase (OD600 0.2) in conditions like altered temperature, agitation, and pH levels, and released phages were quantified. All tested environmental stimuli did not lead to important changes in phage titers from the negative control (NC) titer of roughly 10^6^ to 10^7^ PFU (plaque-forming units)/mL (Fig. 1a).

**Fig. 1:**
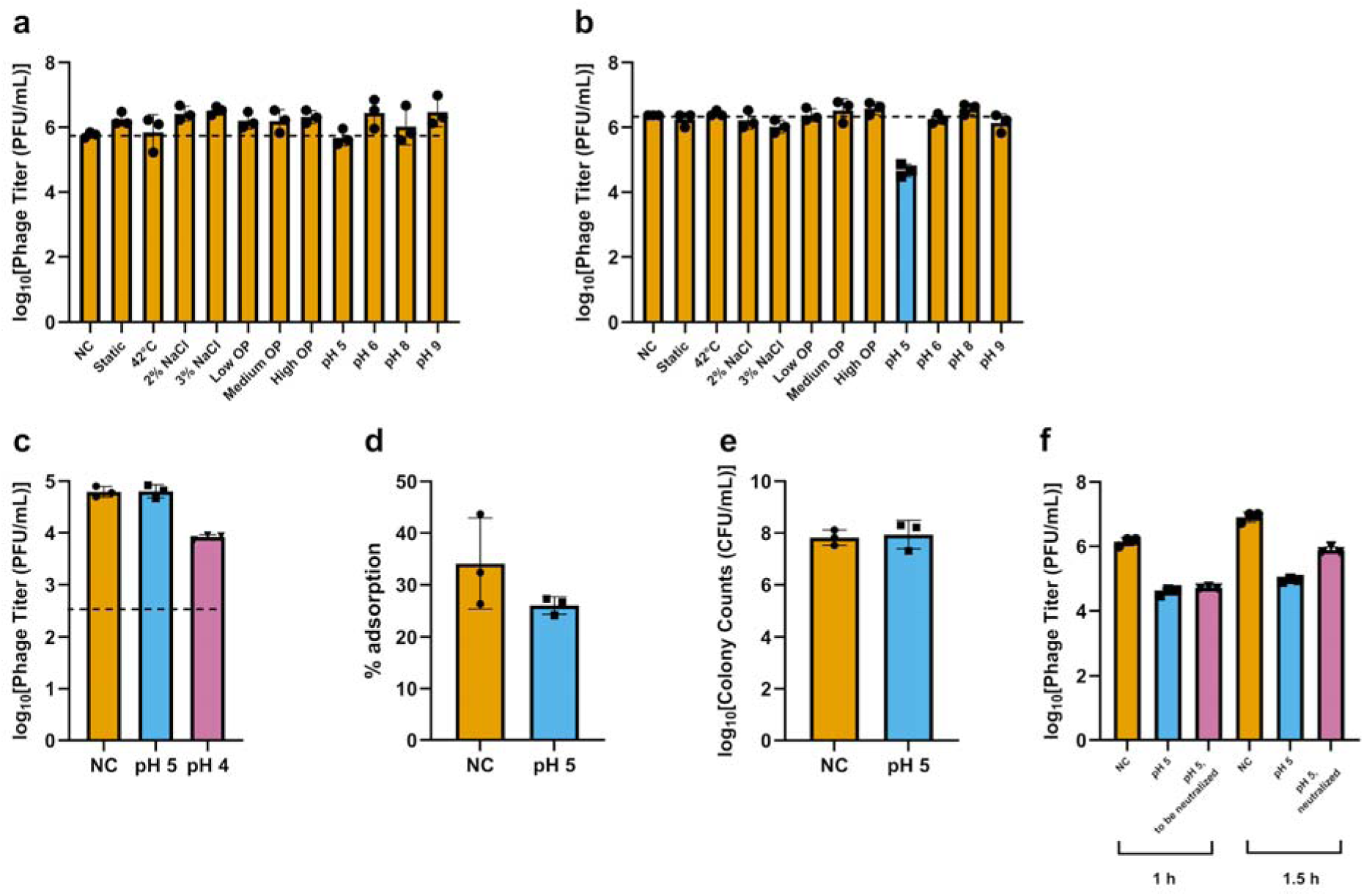
Acute exposure to pH 5 decreases phage titers. a) Chronic exposure to different temperature, agitation, media type, and acidity conditions led to no important changes from baseline in lysogens. Low, medium, and high osmotic pressure (OP) here are achieved using 0.025 g/mL, 0.05 g/mL, and 0.075 g/mL sucrose, respectively. A dashed line is placed at the baseline phage titer. b) Acute exposure to identical conditions as the chronic set generally did not lead to changes in phage titers, except for exposure to a pH of 5, which led to a ∼100-fold reduction. A dashed line is placed at the baseline phage titer. c) Phages remain stable when incubated at pH 5 for 18 h, but degrade at a pH of 4. A dashed line is placed at the lower limit of detection. d) Phage adsorption largely remains stable at pH 5. e) Cells remain viable when incubated at pH 5 for 1 h. f) When cells are exposed to pH 5 for 1 h, and then neutralized to pH 7, they quickly begin to recover within 0.5 h from the suppressive phenotype.

We hypothesized that growing the lysogens in certain environmental conditions was insufficient to evoke a response. Instead, the lysogens and phages may respond better to *changes* in stimuli. We pivoted to using an acute experimental approach. Instead of growing the lysogens to the exponential phase in various conditions, we grew them in optimal conditions (37°C, shaking) to OD600 0.2, and then exposed them for 1 h to the same stimuli as those used in Fig. 1a. This acute condition could better explore any changes in induction as a result of transient host responses.

Many of the conditions did not lead to important changes in the phage titers from the 10^6^ to 10^7^ PFU/mL baseline, except for the acute pH 5 exposure (Fig. 1b), which led to a ∼100-fold decrease in phage titers. Importantly, no such suppression was seen in the chronic pH 5 condition (Fig. 1a). We speculated that perhaps an equivalent change in pH from a different baseline might elicit the same effect. Cells were grown at pH 8 and then brought to pH 6 for 1 h, in a similar manner to the acute condition described in Fig. 1b. No change in titer from baseline was observed (Fig. S1).

Acute exposure to pH 5 specifically is a unique condition that is not replicated by similar changes in pH. The *change* in acidity to pH 5 is necessary for the suppression phenotype.

### The phage titer decrease from acid exposure is likely due to suppression of induction

The low titers caused by the acid exposure does not necessarily mean that induction was being suppressed. Alternative explanations for the drop in titer could be that the phages or cells were not viable or that phages were being more rapidly adsorbed onto cells at that pH, which would deplete measured free phages. We found that the phages were also equally stable when incubated without cells for 18 h at pH 5 or 7. Phages were unstable at pH 4, and this condition led to lower titers (Fig. 1c). This could indicate that suppression of phage production occurs at pHs < 5; we are, however, unable to distinguish this from the degradation that occurs at these pHs.

We tested phage adsorption, or binding to cells, with phages that were incubated with either *E. coli* nonlysogens at pH 5 or 7 (Fig. 1d). The phages largely adsorbed to the same degree in either condition, with a slight trend towards poorer adsorption at pH 5. This trend would yield an increase in free phages, and so does not explain the strong suppression observed. This would suggest an increase in phage titer, if adsorption is impaired, but we instead see suppression.

The lower phage titers could also not be explained by the cells dying; the cells are equally viable between pH 7 and pH 5 conditions after a 1 h incubation, and so the pH 5 condition must involve the same number of cells producing fewer phages (Fig. 1e). They are also capable of recovering from this phenotype, and can still produce phages later; when lysogens are brought to pH 5 for 1 h, and then neutralized back to the pH of control cultures at around pH 7, the titers start to recover only 0.5 h after neutralization (Fig. 1f). These results confirmed that the drop in phage titers after acid exposure was likely due to a suppression of phage production.

### Acid exposure overrides canonical DNA damage-mediated induction

We identified acute acid exposure as a phage suppressor, and sought to investigate whether this phenotype would be potent enough to block phage activity associated with DNA damage.

Mitomycin C (mitC) is a DNA crosslinker.^21^ The DNA damage it causes triggers phage induction in an SOS response-dependent manner, and can drive phage titers up in *E. coli* lambda lysogens from 10^6^-10^7^ PFU/mL up to ∼10^10^ PFU/mL (Fig. 2a). *E. coli* lysogens were grown to exponential phase, and then exposed to either acute pH 5 exposure, mitC, or a combination of the two. pH 5 does not inhibit cell growth, and does not prevent mitC-associated killing over 18 h (Fig. S2). Phage titers were observed at each hour after exposure until 6 h, and also at 18 h (Fig. 2a). Crucially, acid exposure was capable of completely overriding the phage-inducing effect of mitC in the combined condition’s early time points. This suppressive phenotype does not persist for more than a few hours. The decrease in titers tapers off after 4 to 5 h of exposure, and is not detectable at 18 h. Still, this phenotype, however transient, forms an important step in preventing phage induction and toxin production associated with antibiotic use in STEC infections. Acid exposure is able to suppress phage induction, in a way that can override the canonical DNA-damage to induction pathway.

**Fig 2:**
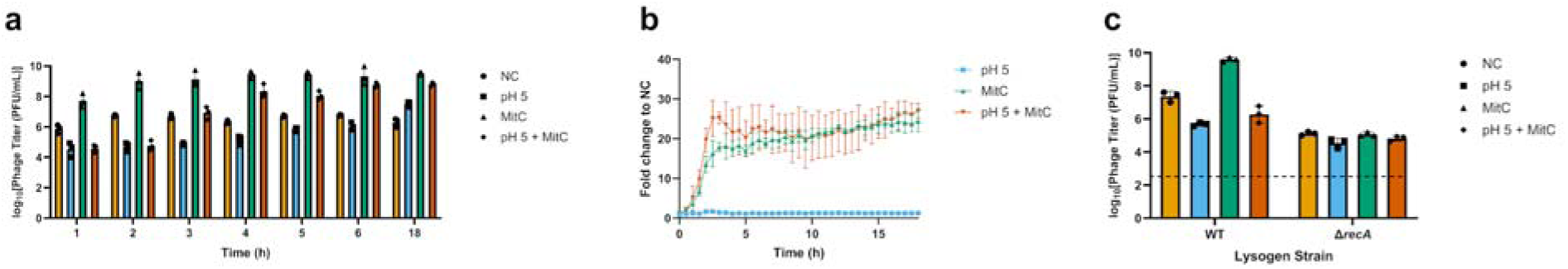
pH 5-associated phage suppression is transient and *recA*-independent. a) Acid exposure transiently reduces phage titers in a manner that overrides mitC (0.5 μg/mL)-associated induction. b) *recA* transcriptional activity in a reporter strain increases with mitC (0.5 μg/mL), independently of whether the cells are at pH 5. c) In a Δ*recA* strain, there is some limited suppression with pH 5. MitC does not lead to any induction in this mutant. A dashed line is placed at the lower limit of detection.

We explored this acid exposure and mitC combination condition more closely through the lens of bacterial SOS responses. When the phage genome goes dormant during lysogeny, cI, a phage protein, represses lytic gene transcription.^1^ The DNA-damaging stimuli discussed earlier trigger the SOS response. The bacterial protein RecA binds to damaged DNA, which causes the bacterial repressor protein LexA to cleave itself, relieving the repression of SOS response genes, including *recA* and *lexA*, and initiating DNA repair.^1^ As the phage repressor protein cI is structurally similar to LexA, it also undergoes cleavage in the presence of the complex. This relieves the repression of lytic genes, inducing the phage.^1^

To examine how this interacts with phage suppression, we exposed an *E. coli* reporter strain for *recA* to either acute pH 5 exposure, mitC at 0.5 μg/mL, or a combination of the two (Fig. 2b). Compared to the negative control, cells at pH 5 did not show *recA* activity; acid exposure does not activate the SOS response. As expected, mitC does increase *recA* transcriptional activity. However, this was entirely unaffected by the presence of acid; acid exposure is able to suppress phage titers despite not interfering with the early steps of the SOS pathway. Additionally, we exposed Δ*recA E. coli* lysogens to the same conditions (Fig. 2c). As expected, this strain does not experience phage induction from mitC. The difference in titer is not as apparent due to the low baseline titres expected from a *recA* mutant, closer to the limit of detection, but there is a 50% reduction between the NC and pH 5 conditions. The effects of acute acid exposure on phage suppression appear *recA*-independent and occur despite SOS activation.

### Trimethoprim acts as a mimic of acute acid exposure in suppressing phage titers

Trimethoprim is a commonly used antibiotic which inhibits the synthesis of folate, an essential precursor for nucleic acid synthesis.^22^ Mitosch et al. found that, through a mechanism that has not been fully elucidated, this leads to the buildup of intracellular protons and an associated acid stress response.^22^ We hypothesized that it could be provide an orthogonal approach to verify the importance of acid stress in phage suppression, and would be more applicable in a clinical setting than acidification.

We exposed *E. coli* lysogens to trimethoprim at the concentrations used by Mitosch et al. (0.5 μg/mL) to elicit the acid stress response, or mitC at 0.5 μg/mL, or a combination of the two (Fig. 3a). This led to the same potent phage suppression phenotype with trimethoprim, in a manner that also overrode the induction associated with mitC. The suppression relative to baseline also persisted for several hours, before it recovered at 18 h. Excitingly, in contrast to pH 5, trimethoprim is able to block induction from mitC even at 18 h. In terms of cell killing, both TMP and mitC kill cells at these concentrations over 18 h (Fig. S3). Unlike mitC, which leads to pronounced induction, TMP does not change phage titers from baseline even at high concentrations at 18 h (Fig. S4).

**Fig. 3:**
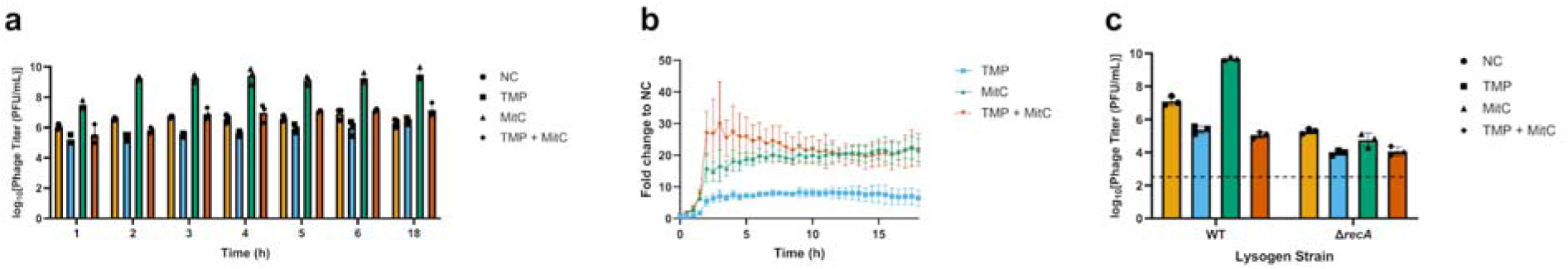
Trimethoprim-associated phage suppression is partially transient and *recA*-independent. a) Trimethoprim (0.5 μg/mL) reduces phage titers in a manner that overrides mitC (0.5 μg/mL)-associated induction. b) *recA* transcriptional activity in a reporter strain increases with mitC (0.5 μg/mL), independently of whether the cells are exposed to trimethoprim (0.5 μg/mL). c) In a Δ*recA* strain, trimethoprim (4 μg/mL) leads to phage suppression and overrides mitC (0.5 μg/mL)-associated induction. A higher trimethoprim concentration was used because the base strain here is *E. coli* BW25113, which requires higher concentrations than K12 for the same suppressive effect. A dashed line is placed at the lower limit of detection.

We also explored trimethoprim in the context of the SOS response. In the *recA E. coli* reporter strain, trimethoprim does lead to *recA* transcriptional activity, likely due to its inhibition of nucleic acid synthesis (Fig. 3b). Similar to pH 5 (Fig. 2b), it also does not alter the *recA* activity caused by mitC. Trimethoprim also behaves very similarly to the pH 5 condition in a Δ*recA E. coli* strain, where its phage suppression is independent of *recA* (Fig. 3c). Trimethoprim acts as an intracellular mimic of the acute pH 5 exposure, with a longer-lived suppressive effect, giving us a better tool to explore this pathway.

### Trimethoprim’s suppressive effect is dampened in acid resistance knockout strains

As both orthogonal approaches led to the same phage suppression phenotype, we explored bacterial responses that could be responsible. *E. coli* harbors several acid stress response systems, many of which are redundant. Of these, the glutamate-dependent acid resistance system (GDAR) is the most potent.^23^ In this system, two isoforms of a glutamate decarboxylase, GadA and GadB, convert glutamate to GABA and consume protons in the process, reducing intracellular acidity. GadC, a membrane protein, transports the GABA out while taking more glutamate into the cell for the decarboxylation to continue.^24^ Interestingly, GadB is located in the cytoplasm at pH > 5.6, and below 5.6 it colocalizes with the cellular membrane fraction where it is more efficient.^23^ GadC also only changes its conformation and activates at pH ≤ 5.5.^23^ This may align well with our observations related to pH 5 being a unique condition.

Knockout strains of *gadA*, *gadB*, and *gadC* did not show differences in their baseline phage production compared to wild-type (Fig. 4a), but did show an increased sensitivity to extremes in pH. Δ*gadC* strains were more likely to die when brought to pH 3, but trimethoprim and pH 5 acid exposure did not appear to have an effect on viability (Fig. 4b).

**Fig. 4:**
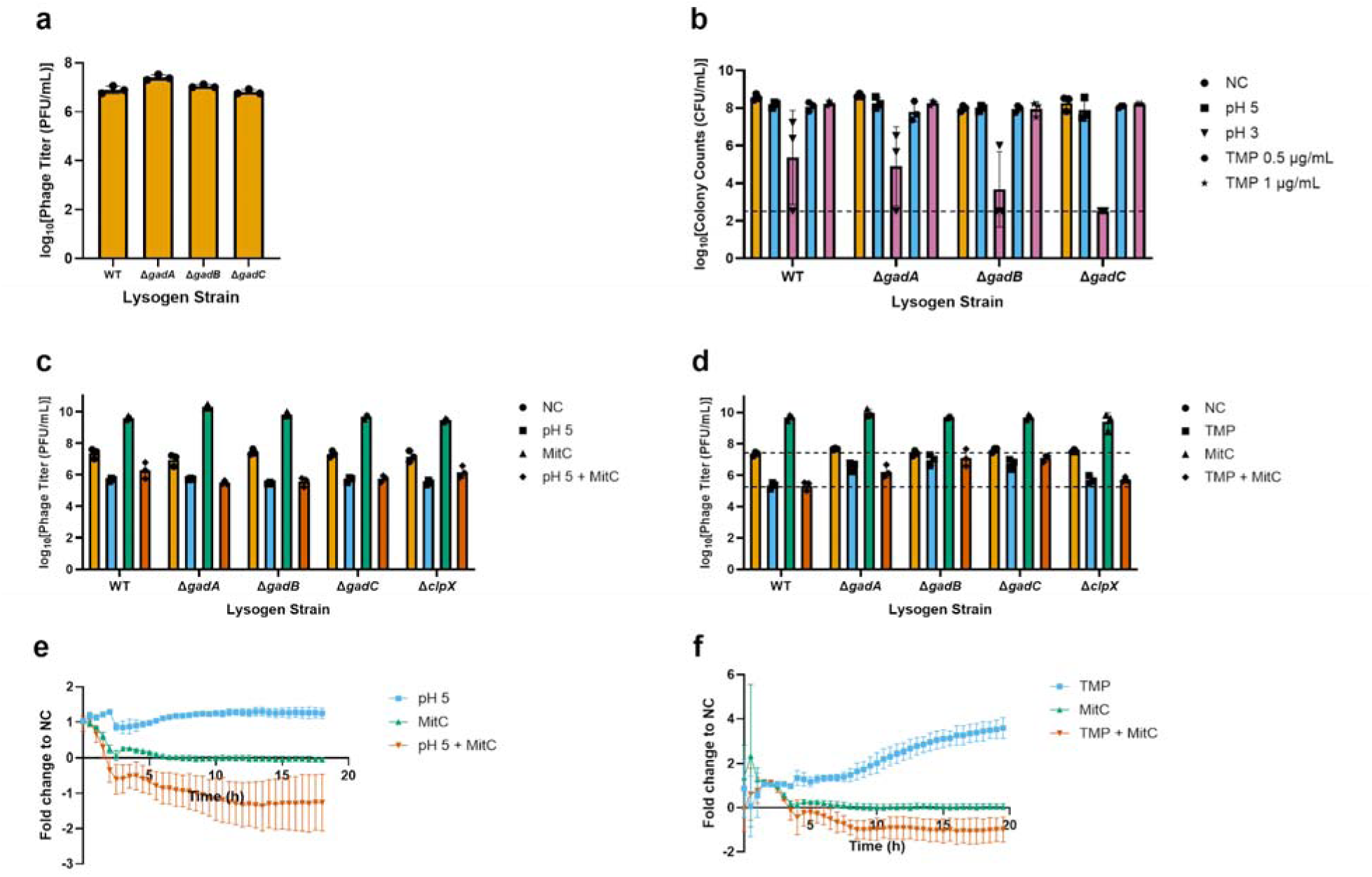
Trimethoprim, but not pH 5 exposure, appears to partially depend on the GDAR to suppress phages. a) GDAR knockout strains show no change in baseline phage titer compared to a wild-type strain. b) GDAR knockout strains show poorer survival at pH 3 compared to a wild-type strain. c) pH 5 suppresses phages in a manner that overrides mitC (0.5 μg/mL)-associated induction even in the GDAR and *clpX* mutants. d) Trimethoprim (4 μg/mL) still overrides mitC (0.5 μg/mL)-associated induction in the knockout strains, but suppression relative to the NC is not as pronounced. Dashed lines are placed to show baseline phage titers and suppression in the wild-type strain. e) pH 5 does not lead to a pronounced increase in *gadBC* transcriptional activity, but mitC (0.5 μg/mL) suppresses it. f) Trimethoprim (0.5 μg/mL) increases *gadBC* transcriptional activity, but mitC (0.5 μg/mL) suppresses it.

The suppressive effects of pH 5 exposure, and its ability to override mitC-associated induction, happen independently of whether various genes involved in the GDAR are knocked out (Fig. 4c); this could mean that the cell is still activating the acid stress response through other pathways. Downstream to the GDAR, the RpoS general stress response is activated, which is known to be linked to decreased Stx phage induction.^18^ RpoS accumulates under stress, but is rapidly degraded by ClpXP protease during normal growth.^25^ We tested a Δ*clpX* strain, and it also showed no difference in phage suppression from pH 5 compared to wild-type.

The same observations were not seen with trimethoprim exposure. In knockout strains of *gadA*, *gadB*, and *gadC*, trimethoprim showed dampened suppression compared to baseline, even though its ability to override mitC-associated induction remained. The Δ*clpX* strain showed no difference in phage suppression from trimethoprim compared to wild-type (Fig. 4d). These findings align well with what we find in a *gadBC* reporter strain. pH 5 exposure does not lead to changes in *gadBC* transcriptional activity, while trimethoprim leads to a mild increase (Fig 4e-f). pH 5 can suppress phages even in the absence of the GDAR, while trimethoprim partially relies on this pathway.

Interestingly, mitC leads to a dampening of *gadBC* activity, independent of whether pH 5 or trimethoprim are present (Fig 4e-f), showing a connection between the GDAR and DNA damage. This relationship has been explored to some extent in the literature. Mitosch et al. found that trimethoprim depletes adenine nucleotides in the cell, and decreases intracellular ATP levels.^22,26^ ATP biosynthesis is required for both acid resistance and DNA-damage responses in *E. coli*.^27^ This may help explain our findings in Fig. 4e-f, and additionally connects to how apyrase, which cleaves ATP, is known to also reduce phage induction and Stx production.^16^

### Trimethoprim allows for the activation of the SOS response and lytic induction genes, but suppresses the amount of non-integrated, circularized phage DNA

After exploring the bacterial side of SOS and acid stress responses involved in suppression, we moved to investigating how trimethoprim affects phage activity and replication. To do this, we first collected lysogen samples every 20 min for 2 h, and at 4 h post-exposure to trimethoprim, mitC, or a combination of the two, where we observed phage suppression with greater early timepoint resolution (Fig. 5a). Then, we used RT-qPCR to assess the transcriptional activity of genes in these samples that link the SOS response to phage induction by quantifying their mRNA levels. *recA* and *lexA* mRNA levels were elevated when both trimethoprim and mitC were present (Fig. 5b-c). This matches the results in Fig. 3b, where both substances still lead to SOS activation. Again, this shows us that phage suppression must occur independent of this effect. *cro* is a key lytic induction gene; it is transcribed when cI repression is relieved, and in turn, shuts down *cI* transcription and inhibits lysogeny.^28^ We found that *cro* mRNA levels are increased when both trimethoprim and mitC were present (Fig. 5d), which means that phage induction is starting when trimethoprim is present. Interestingly, *cI* mRNA levels are only elevated when mitC is present, and are suppressed in the combination condition (Fig. 5e).

**Fig. 5:**
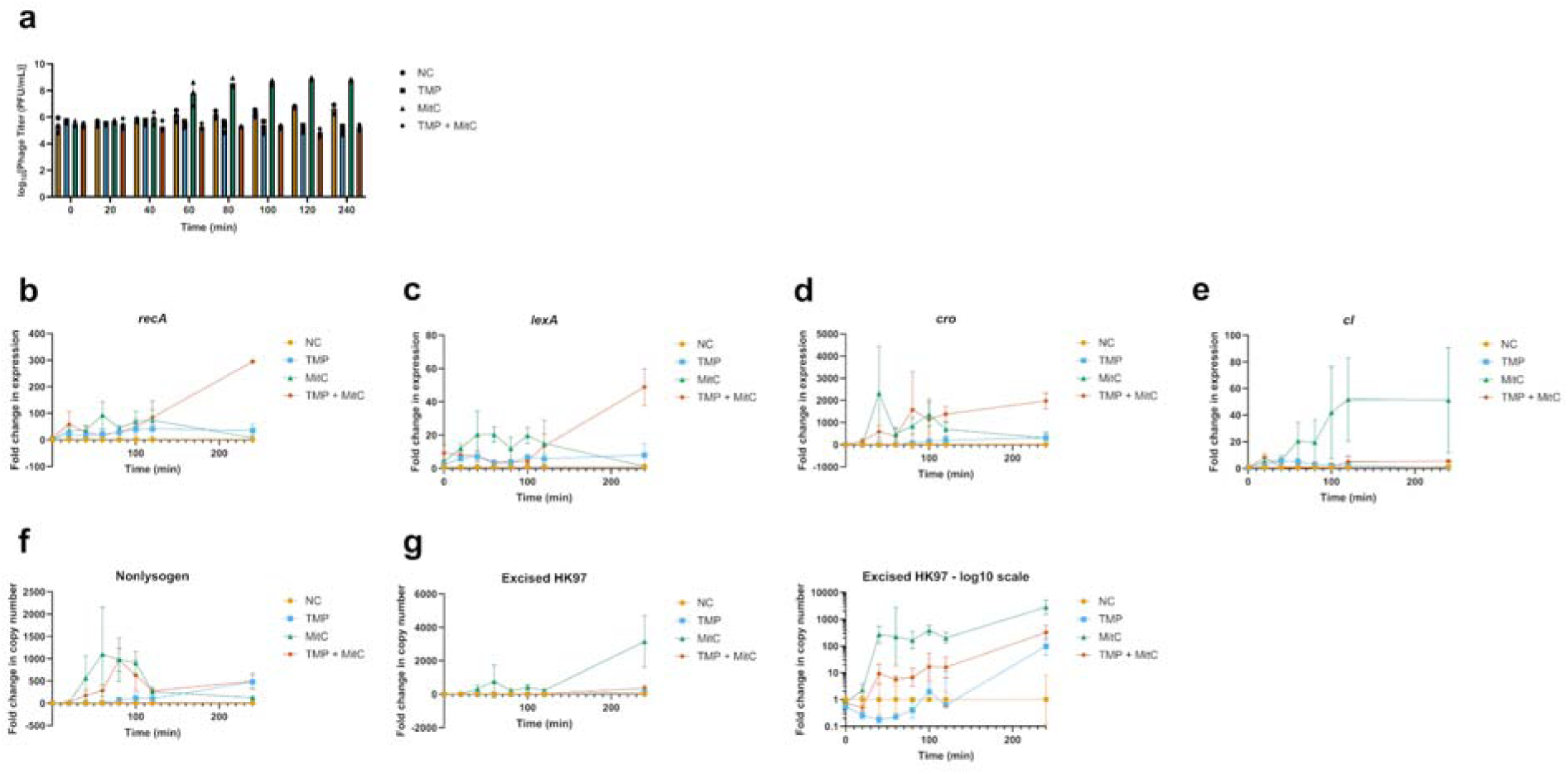
Trimethoprim suppresses phage genome replication. a) With more frequent sampling, we continue to see the trimethoprim (4 μg/mL)-associated suppression that blocks mitC (4 μg/mL)-associated induction. b-d) *recA*, *lexA*, and *cro* transcriptional activity are all increased in the trimethoprim and mitC combination condition. e) *cI* transcriptional activity increases with mitC, but appears to be suppressed when trimethoprim is also present. f) There appears to be an increase in the amount of bacterial DNA where the HK97 genome has excised when cells are exposed to mitC, independently of whether trimethoprim is present. g) The number of non-integrated, circularized HK97 genomes is increased with mitC exposure, but suppressed by trimethoprim. Data are also charted on a log-scale to the right for easier parsing.

According to existing literature, *cro* is known to shut down cI production during induction to prevent lysogeny^28^, which is inconsistent with our findings. Once Cro production is sufficiently high, the phage will switch to lytic replication and levels of cI present will not matter, so while this disconnect is intriguing, it is not clear how it would tie to our suppressive phenotype.

Nevertheless, we can see that the bacterial SOS response and key early lytic induction genes are being activated, even when trimethoprim suppresses overall titers.

We then moved downstream in the phage induction pathway to characterize the specific point in the phage replication cycle where suppression is taking place. Using the same samples collected in Fig 5a, we measured the amount of bacterial DNA showing an empty site where the HK97 prophage has removed itself, as a proxy for the amount of excision happening. This process also forms an early step in phage induction. We see an increase here with both mitC and the combination condition, and even trimethoprim along to a lesser extent (Fig. 5f). Trimethoprim is not blocking induction at the level of preventing excision.

We next measured the amount of non-integrated, circularized HK97 DNA. This is phage DNA that has been removed from the bacterial chromosome, and will either initiate replication or is the result of that replication. MitC alone leads to a large increase in circularized HK97 DNA, while this is suppressed in the combination condition (Fig. 5g). Trimethoprim appears to be allowing for the SOS-mediated de-repression of the phage lysogenic cycle, leading to excision of the prophage from the bacterial chromosome, but not replication of that genome. Importantly, in the case of Stx phages, this would mean that the production of Shiga toxins, occurring during late lytic replication, would be blocked by trimethoprim. Note that this could also mean trimethoprim could simply block phage genome replication purely in the lytic cycle, without requiring lysogeny, which would mean any events where Stx phages attempt to replicate in other, nonlysogen hosts would also be blocked.

### Trimethoprim suppresses Shiga toxin levels in a clinical STEC strain

After characterizing phage suppression associated with acid exposure and trimethoprim in nonpathogenic, laboratory strains, we moved towards examining whether this phenotype translates to clinical, STEC strains.

We exposed a clinical isolate of *E. coli* O103:H25 to trimethoprim, ciprofloxacin (a DNA-damaging antibiotic, which we felt was more clinically relevant than using mitC), or a combination of the two for 2 h and 18 h, and quantified toxin levels using a clinical diagnostic-use ELISA assay (Fig. 6). Toxin production was limited at 2 h and differences between groups were more difficult to parse. However, it was greatly elevated with ciprofloxacin exposure at 18 h, and highly suppressed by trimethoprim. In conclusion, in a clinical STEC strain, we see an overriding suppression of toxin production despite the presence of a DNA-damaging antibiotic. This represents a promising lead towards the goal of treating STEC infections and allowing for cell killing without the risk of phage-associated virulence and HUS.

**Fig. 6:**
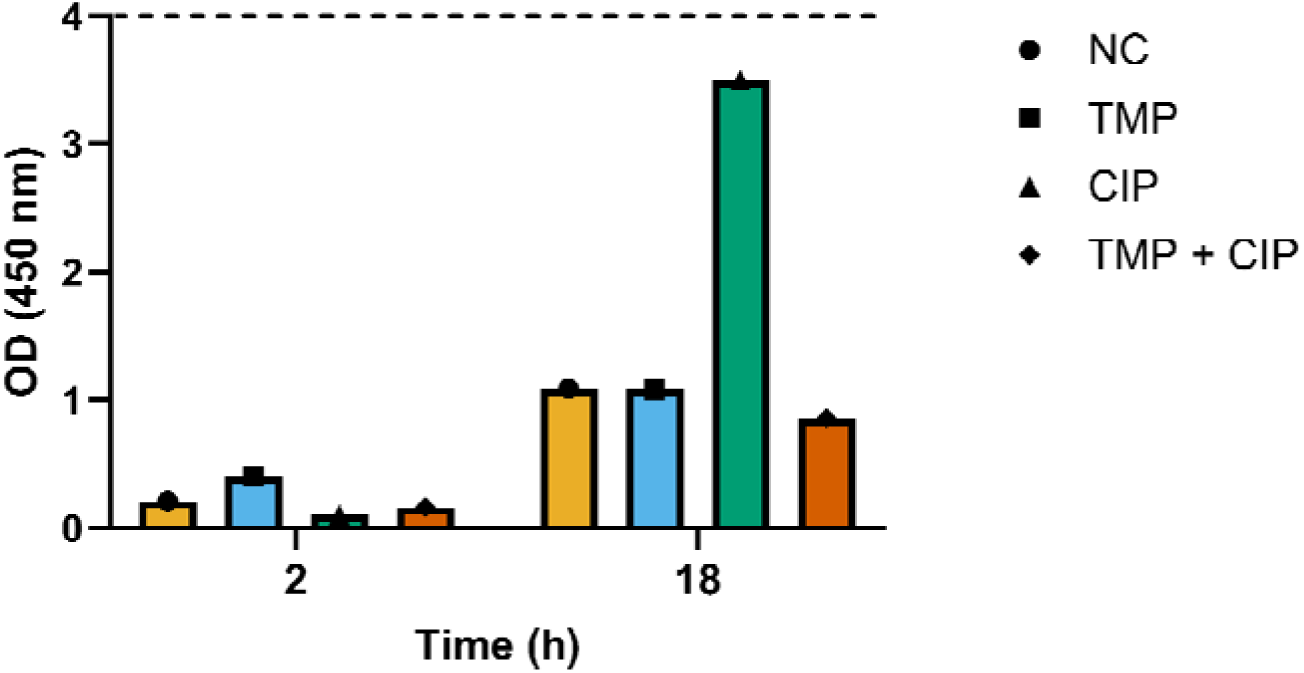
An STEC clinical strain shows a clear overriding suppression of toxin production when exposed to trimethoprim alongside ciprofloxacin. Differences in toxin production are more difficult when cells are tested 2 h post-exposure, because levels are quite low. At 18 h, the clinical strain recapitulates our phage suppression phenotype, and shows a pronounced increase in toxin production when exposed to ciprofloxacin, which is suppressed when trimethoprim is also present. Trimethoprim and ciprofloxacin concentrations here are at 2x MIC. The ELISA readout is considered positive at OD450 ≥ 0.180, indicating the presence of at least 7 pg of Stx1 or 15 pg of Stx2. The dashed line indicates the upper limit of detection.

## Conclusions and future directions

We describe acute acid exposure and trimethoprim exposure as a mechanism for phage suppression, even in the presence of DNA-damaging drugs. This effect is *recA*-independent, and in the case of trimethoprim, is partially dependent on the GDAR response. We found that trimethoprim allows for activation of bacterial SOS responses through increased *recA* and *lexA* transcriptional activity, and also allows for excision of the prophage during induction. It does not block *cro* transcriptional activity, showing that the earliest stages of induction are uninhibited. Instead, it appears to completely block replication of the phage genome.

The acid stress response appears to transiently pause phage production. When examining *E. coli* in the context of its natural environment, the gastrointestinal tract^24^, this phenotype makes sense. Passing through very low pH stomach acid is something wild-type bacteria only experience for a few hours before they pass through to the relatively-neutral pH of the small intestines.

Responding to acid stress may be a short-term phenomenon, allowing phages to avoid being released into a hostile environment where they degrade. TMP, then, is able to intracellularly mimic acid stress and maintain suppression for longer.

We then observed how this suppressive phenotype translates into a clinical STEC isolate, as reduced toxin production even in the presence of ciprofloxacin, a DNA-damaging antibiotic. We plan to continue to explore this phenotype for its clinical use in treating STEC infections, potentially through increased *in vitro* testing and *in vivo* models. We also plan to further characterize the effect in terms of a specific, mechanistic connection between acid stress responses and phage induction.

## Methods

All strains used in experiments are given in Table 1.

**Table 1:** Strains used in this study.

| Isolates/strains | Use | Source |
| --- | --- | --- |
| *Escherichia coli* K-12 Ymel mel-1 supF58 | Lysogenized with HK97. | Dharmacon Keio collection through Horizon Discovery. |
| *Escherichia coli* K-12 BW25113 lysogen HK97 | Base strain for Keio collection<sup>29</sup>, lysogenized with HK97. | Keio collection, Dr. Eric Brown, McMaster University. |
| *Escherichia coli* K-12 BW25113 $\Delta\text{recA}$ lysogen HK97 | Keio collection knockout mutant, lysogenized with HK97. | Dharmacon Keio collection through Horizon Discovery. |
| *Escherichia coli* K-12<br>BW25113 $\Delta gadA$ lysogen<br>HK97 | Keio collection knockout<br>mutant, lysogenized with<br>HK97. | Keio collection, Dr. Eric<br>Brown, McMaster University. |
| *Escherichia coli* K-12<br>BW25113 $\Delta gadB$ lysogen<br>HK97 | Keio collection knockout<br>mutant, lysogenized with<br>HK97. | Keio collection, Dr. Eric<br>Brown, McMaster University. |
| *Escherichia coli* K-12<br>BW25113 $\Delta gadC$ lysogen<br>HK97 | Keio collection knockout<br>mutant, lysogenized with<br>HK97. | Keio collection, Dr. Eric<br>Brown, McMaster University. |
| *Escherichia coli* K-12<br>BW25113 $\Delta clx$ lysogen<br>HK97 | Keio collection knockout<br>mutant, lysogenized with<br>HK97. | Keio collection, Dr. Eric<br>Brown, McMaster University. |
| *Escherichia coli* K-12<br>MG1655 pUA66 empty<br>vector control | Base strain for Alon library.<sup>30</sup> | Alon library, Dr. Eric Brown,<br>McMaster University. |
| *Escherichia coli* K-12<br>MG1655 pUA66-*recA*<br>reporter | Alon library promoter-GFP<br>reporter for *recA*. | Alon library, Dr. Eric Brown,<br>McMaster University |
| *Escherichia coli* K-12<br>MG1655 pUA66-*gadBC*<br>reporter | Alon library promoter-GFP<br>reporter for *gadBC*. | Alon library, Dr. Eric Brown,<br>McMaster University. |
| *Escherichia coli* O103:H25 | Clinical STEC strain. | Dr. Marek Smieja, Hamilton<br>Health Sciences. |

### Chronic versus acute shock challenge

100 μL of overnight *E. coli* K12 HK97 lysogen bacterial culture was subinoculated into 10 mL LB. In the chronic exposure condition, the temperature, agitation, and other factors could be manipulated, and the bacteria were allowed to grow to an OD600 of 0.2. pH levels were adjusted using the addition of 6 M HCl or 2 M NaOH. A 1 mL sample was taken and filtered using a 0.45 μm filter, and phage titers were quantified using a plaque assay. For the acute exposure condition, the subinoculated culture was grown to an OD600 of 0.2 in a 37°C, 250 RPM shaking incubator, and then shocked with the desired condition for 1 hour. Again, a 1 mL sample was filtered and phage titers were quantified using a plaque assay.

### Plaque assay

10-fold serial dilutions of a desired phage sample were prepared in LB, down to a dilution factor of 10^7^. 300 μL of overnight bacterial culture that is susceptible to the phage was mixed with 3 mL 0.75% LB soft agar at 55°C, and then poured onto a 1% LB agar plate and allowed to dry. 3 μL of each phage dilution was placed onto the plate, allowed to dry, and the plate was then allowed to incubate at 37°C overnight. Plaques could then be counted.

### Colony-forming unit assay

10-fold serial dilutions of a desired bacterial sample was prepared in LB, down to a dilution factor of 10^7^. 3 μL of each dilution was placed onto a 1% LB agar plate, allowed to dry, and the plate was then allowed to incubate at 37°C overnight. Colonies could then be counted.

### Knockout strains

Overnight *E. coli* Δ*recA*, Δ*gadA*, Δ*gadB*, and Δ*gadC* HK97 lysogen strains from the Keio were grown in 50 μg/ml kanamycin. 100 μL of each was subinoculated into 10 mL LB, and grown to an OD600 of 0.2 in a 37°C, 250 RPM shaking incubator, before subsequent exposures to antibiotics or pH levels were carried out. In these experiments, the WT control was *E. coli* BW25113, the base strain used to generate these knockouts, lysogenized with HK97.

### Fluorescence assay

Overnight *E. coli* recA GFP, gadBC GFP, and empty vector control reporter strains from the Alon library^30^ were grown in 50 μg/ml kanamycin. 100 μL of each was subinoculated into 10 mL LB, and grown to an OD600 of 0.2 in a 37°C, 250 RPM shaking incubator. A clear black-bottom 96-well plate with desired pH and antibiotic levels was prepared. 100 μL of each reporter culture was placed in assigned sample wells, and each well was filled to a total volume of 250 μL with sterile distilled H_2_O. The assay was then placed in a plate reader with the following settings: Nunc 96-well optical bottom plate with lid, 37°C, 1°C gradient, start kinetic (runtime 18:00:00, reads 37 (shake double orbital (00:05), read absorbance 600 nm, read fluorescence intensity 479, 520), end kinetic. Raw fluorescence values were blanked against the empty vector control strains and normalized to OD600 to account for cell growth. Fold changes in fluorescence intensity were expressed relative to negative controls.

### qPCR

Genomic DNA extraction was done on all samples using an NEB Monarch Spin Genomic DNA Purification Kit. DNA concentrations were quantified using a high-sensitivity dsDNA Qubit fluorometer assay. Each sample was amplified using primers designed to detect the phage-host junction with integrated HK97, without integrated HK97, and circularized, excised HK97 copies, along with housekeeping gene *cysG*. PowerUp SYBR Green Master Mix was used to conduct the qPCR, with the following settings: initial denaturation at 94°C for 2 minutes, then 40 cycles of denaturation at 95°C for 15 seconds and annealing/extension at 60°C for 1 minute. A melt curve was also generated from 65°C to 95°C in 0.5°C increments per second. Data were analysed using the Pfaffl method^31^, normalized to negative control averages and housekeeping gene levels to give a fold change in copy number. All primers used are given in table 2.

**Table 2:**
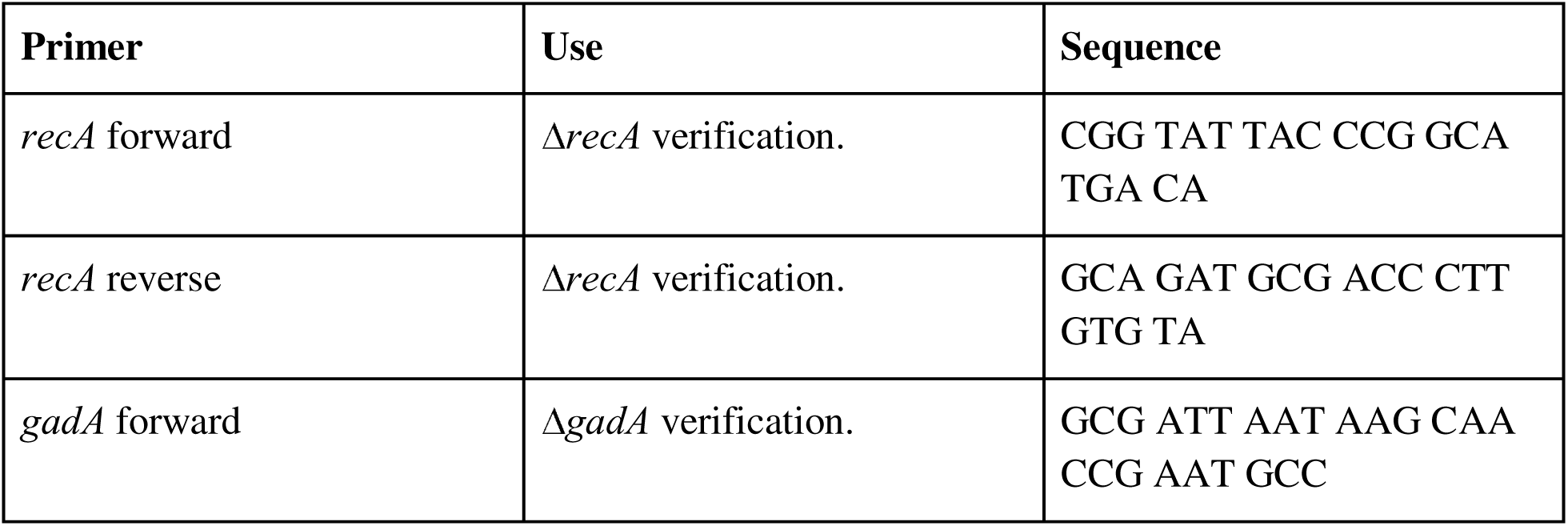
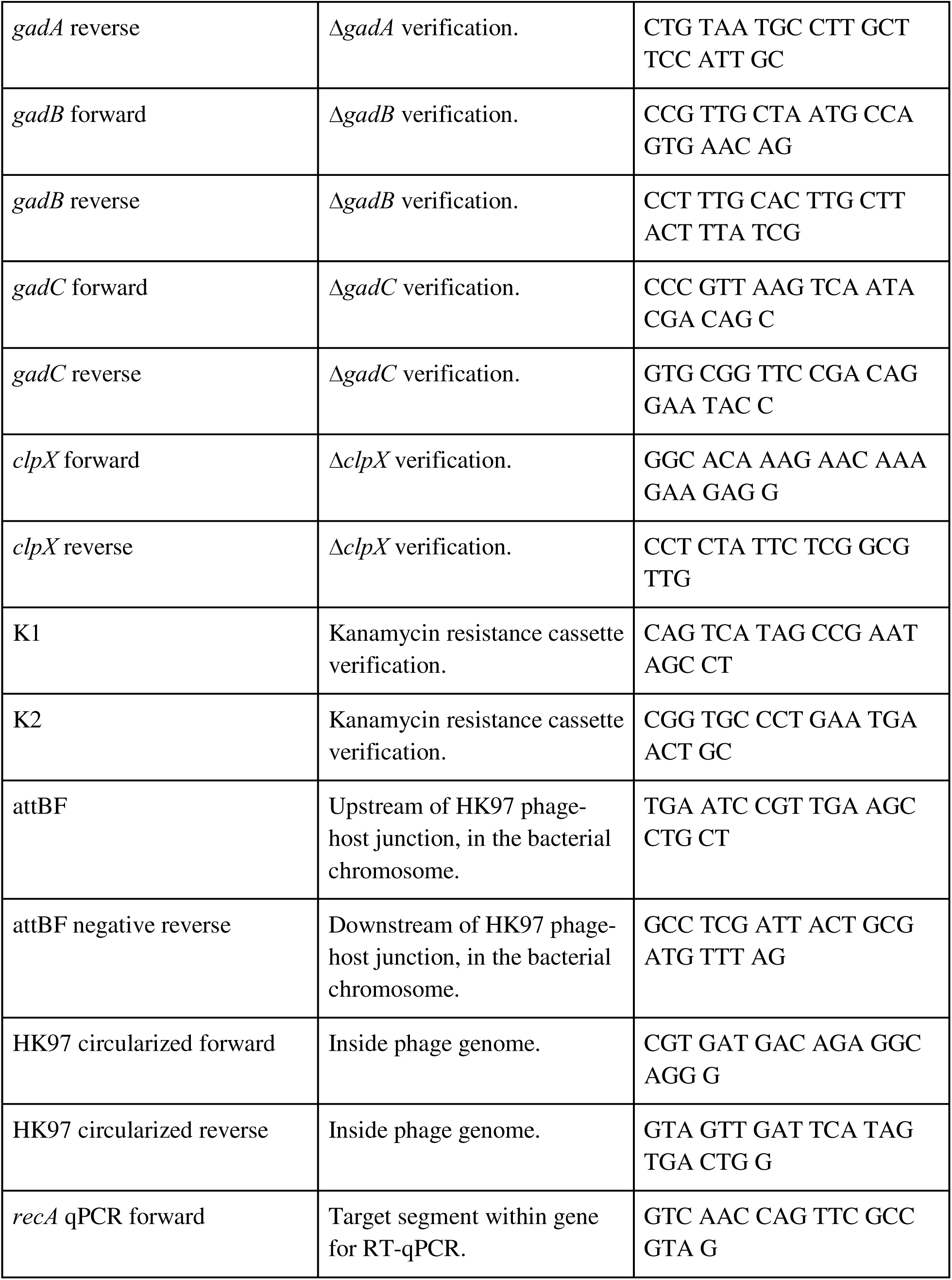
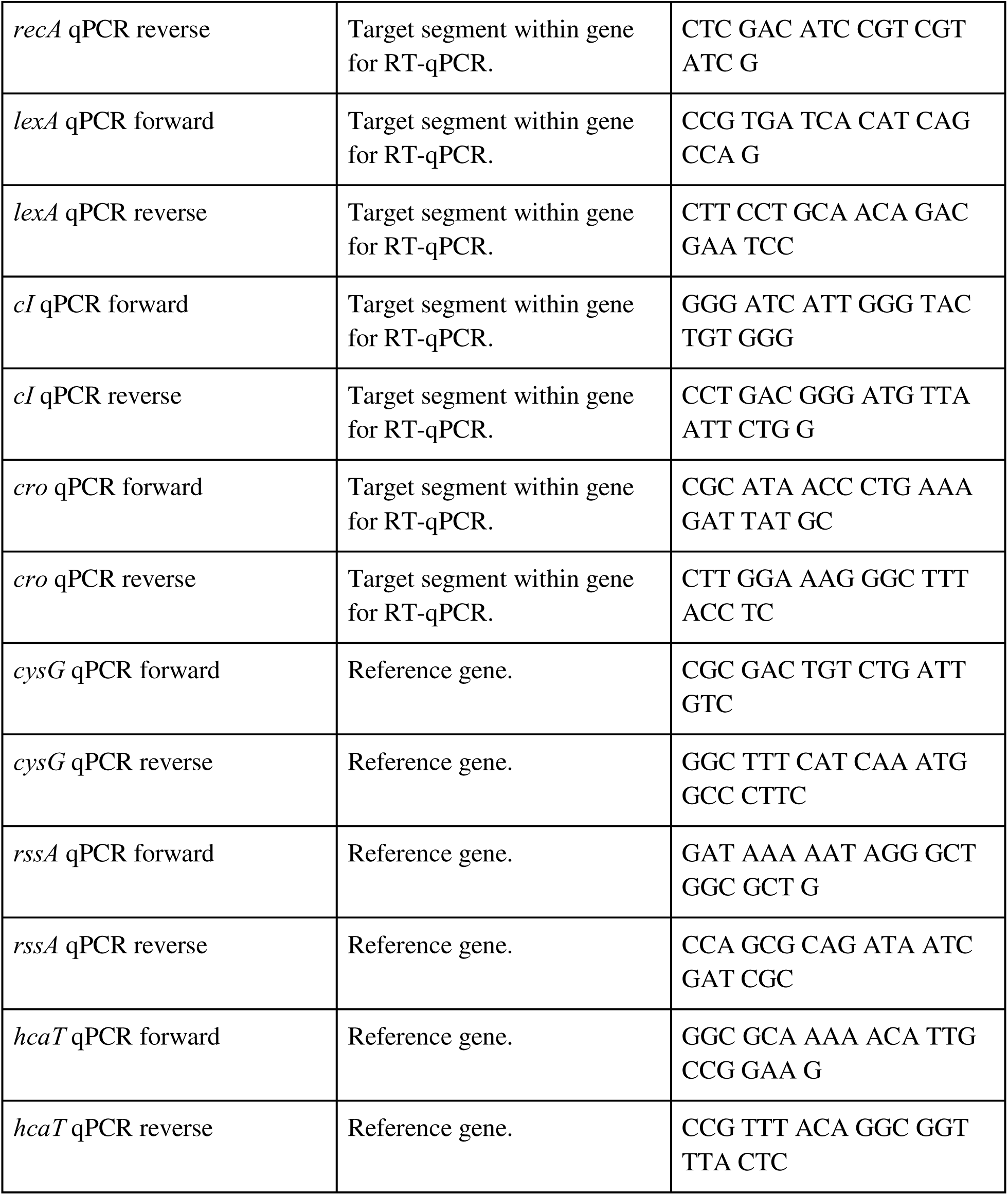
Primers used in this study.

### RT-qPCR

RNA extraction was done on all samples using an NEB Monarch Spin RNA Isolation Kit. An RNA gel was conducted to verify the presence of 23S and 16S ribosomal RNA, and concentrations were quantified using a high-sensitivity RNA Qubit fluorometer assay. A Bio-Rad iScript cDNA Synthesis Kit was used to convert all samples to cDNA. Each sample was amplified using primers designed to detect *recA*, *lexA*, *cI*, and *cro* cDNA, along with housekeeping genes *cysG*, *rssA*, and *hcaT*. PowerUp SYBR Green Master Mix was used to conduct the qPCR, with the following settings: initial denaturation at 94°C for 2 minutes, then 40 cycles of denaturation at 95°C for 15 seconds and annealing/extension at 60°C for 1 minute. A melt curve was also generated from 65°C to 95°C in 0.5°C increments per second. Data were analysed using the Pfaffl^31^ method, normalized to negative control averages and housekeeping gene levels to give a fold change in expression. All primers used are given in table 2.

### STEC ELISA

Clinical STEC isolate colonies were picked and incubated in 200 μL LB overnight. 2.5 μL of these overnight cultures were subinoculated into a total volume of 250 μL of LB, and grown to OD600 0.2. 96-well plates with desired antibiotic levels were prepared. 100 μL of each STEC culture was placed in assigned sample wells, and each well was filled to a total volume of 250 μL with sterile distilled H_2_O. These were allowed to grow for 2 or 18 h, and then filtered using 0.45 μm filter plates. A Meridian Premier EHEC microwell enzyme immunoassay for the detection of Stx1 and Stx2 was used to analyse toxin levels in the filtrates, with readouts at an absorbance of 450 nm taken with a plate reader.

---

**Fig. S1:**
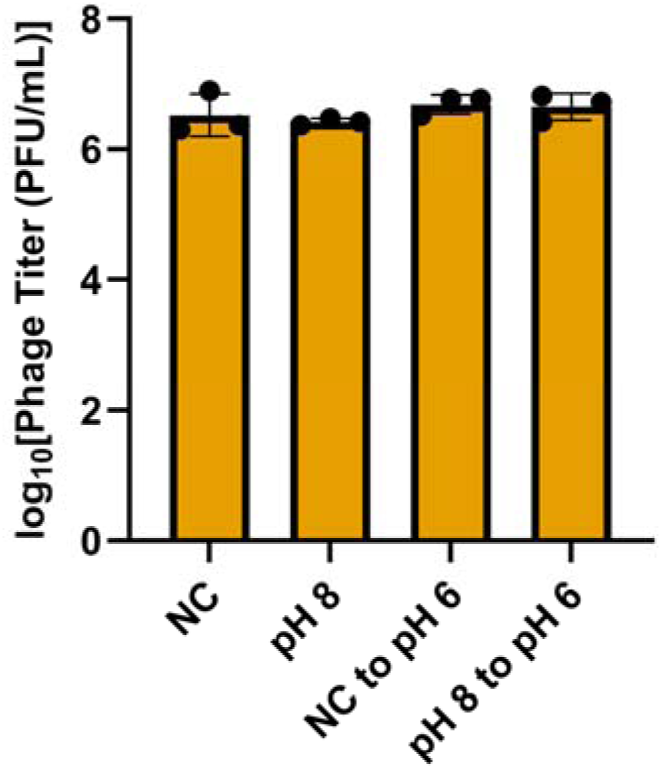
The reduction in titers from acute acid exposure to pH 5 is not replicated by growing cells in pH 8 broth, and then bringing them to a pH of 6.

**Fig. S2:**
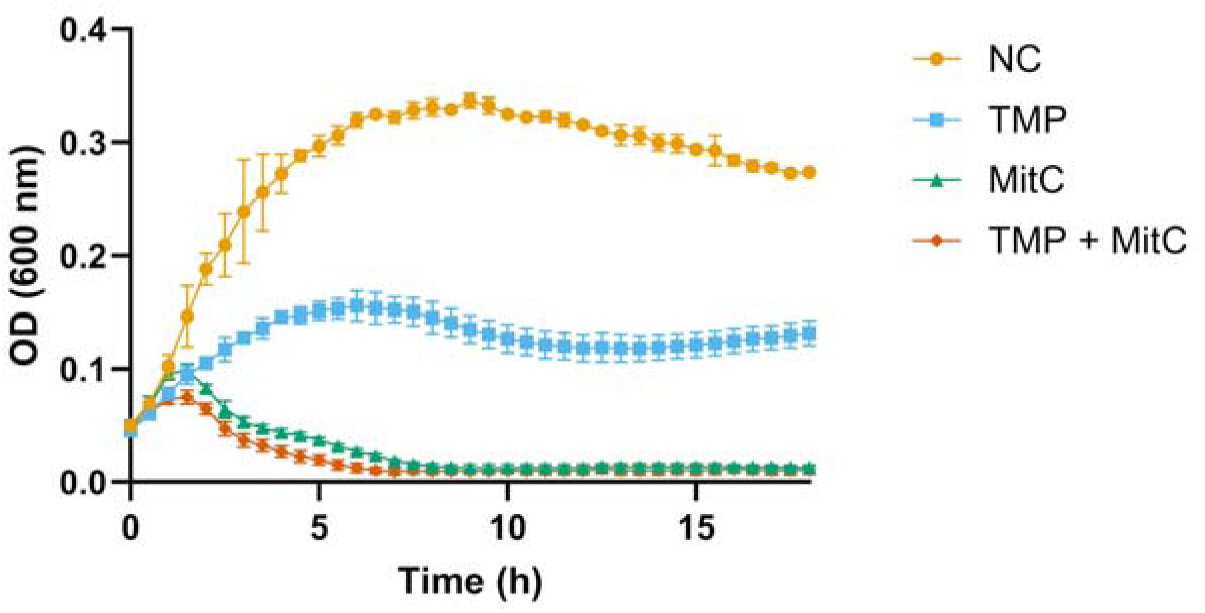
pH 5 causes a small growth delay, but reaches the same final OD600 as NC at 18 h. pH 5 does not prevent mitC (0.5 μg/mL)-associated killing.

**Fig. S3:**
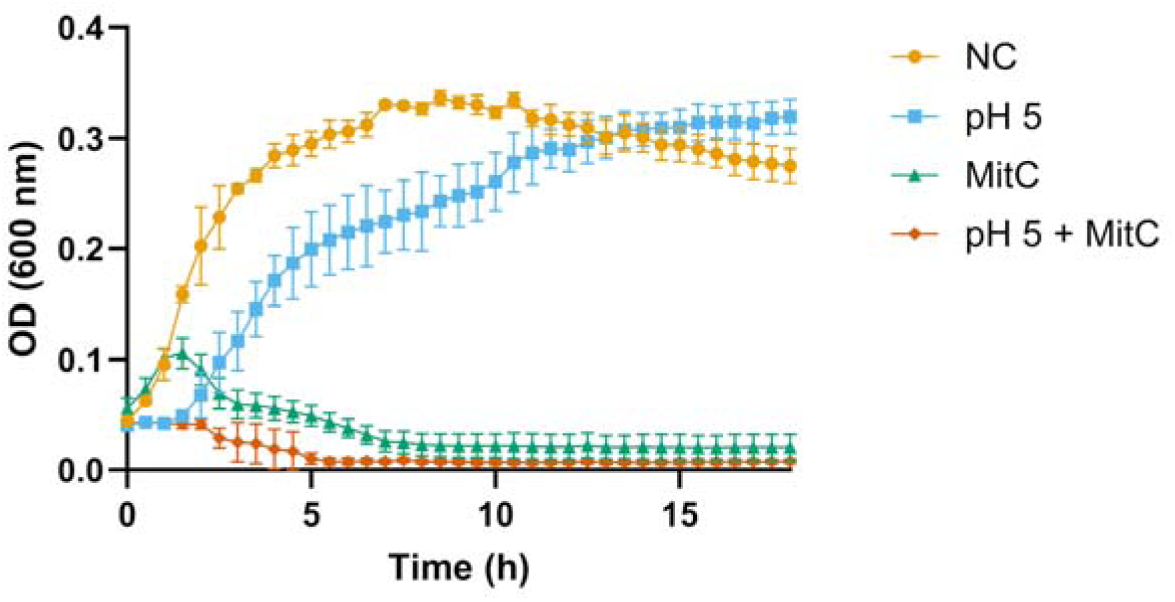
TMP (0.5 μg/mL) and mitC (0.5 μg/mL) both lead to cell death over an 18 h growth curve, and synergize in killing in the combination condition.

**Fig. S4:**
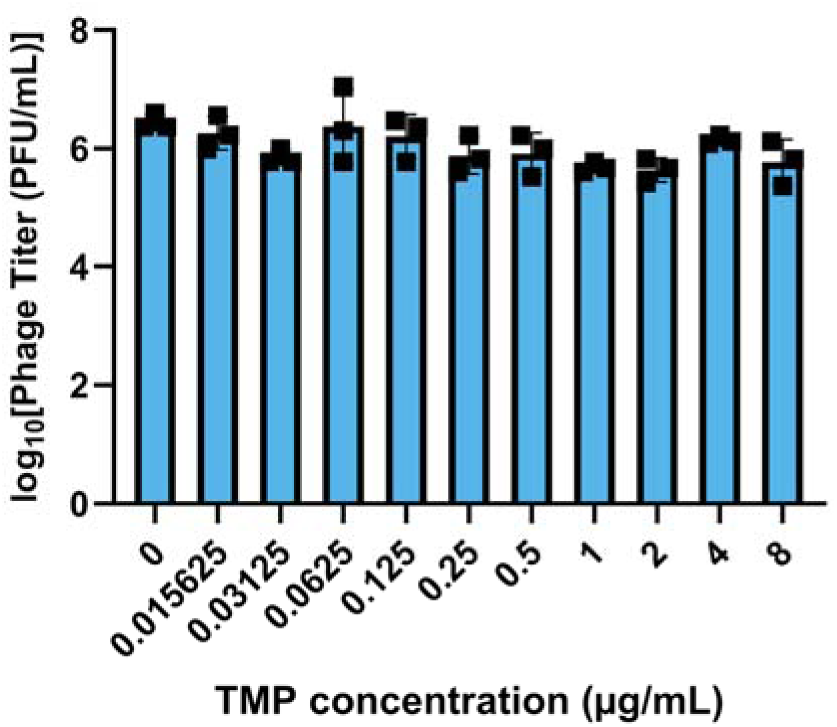
TMP, even at high concentrations, does not cause an important change in phage titers at 18 h. MIC here is around 0.5 to 1 μg/mL.

